# A Suv39H1-low chromatin state drives migratory cell populations in cervical cancers

**DOI:** 10.1101/241398

**Authors:** Calvin Rodrigues, Chitra Pattabiraman, Suma Mysore Narayana, Rekha V. Kumar, Dimple Notani, Patrick Varga-Weisz, Sudhir Krishna

**Affiliations:** National Centre for Biological Sciences, Tata Institute of Fundamental Research, GKVK-UAS, Bangalore, India; Department of Pathology, Kidwai Cancer Institute, Bangalore, India; The Babraham Institute, Cambridge, UK; School of Biological Sciences, University of Essex, Colchester, UK

## Abstract

The emergence of migratory cell populations within tumours represents a critical early stage during cancer metastasis. We have previously reported one such population, marked by CD66, in cervical cancers. It is unclear what broad mechanisms regulate such migratory populations. Here, we describe the role of a Suv39H1-low heterochromatin state as a driver of cervical cancer migratory populations. Cervical cancer cells sorted based on migratory ability *in vitro* show low Suv39H1, and Suv39H1 knockdown enhances cell migration. Histopathology shows the emergence of migratory Suv39H1^low^ populations in advanced carcinoma progression. Meta-analysis of data from The Cancer Genome Atlas (TCGA) reveals that Suv39H1-low tumours show migration and CD66 expression signatures, and correlate with lower patient survival. Lastly, genome-wide profiling of migrated populations using RNA-Seq and H3K9me3 ChIP-Seq reveals Suv39H1-linked transcriptome alterations and a broad loss of H3K9me3, suggesting an increase in chromatin plasticity in migrated populations. The understanding of such chromatin based regulation in migratory populations may prove valuable in efforts to develop anti-metastatic strategies.

## Introduction

Despite significant advances in vaccination and screening, cervical cancer represents one of the most common cancers in women in the developing world. A large incidence of cancer associated mortality is due to metastatic progression, and the emergence of migratory subpopulations within tumours represents a rate limiting step during this process. We have previously reported that one such population in cervical cancers, marked by CD66^high^, shows enhanced migratory potential, gene expression signatures of migration, and associations with metastasis^1–3^. However, it is not clear what broad mechanisms regulate such migratory populations, and how these cells would retain a “memory” of initial migration-inducing cues during metastasis. It is also unknown how such mechanisms would permit plasticity, the ability to switch between migratory and proliferative states, allowing the formation of macroscopic distant metastases.

We have previously reported that CD66-enriched spheroid cultures show a distinct transcriptional profile and altered expression of several chromatin regulators, including a depletion of heterochromatin regulator Suv39H1^1,2^. Heterochromatin based modulation would allow plasticity and memory in migratory populations, through metastable changes in chromatin conformation. Further, Suv39H1-regulated heterochromatin has been demonstrated to play a barrier to cell fate transitions during IPSC reprogramming and T-cell lineage commitment^4–8^. In this study, we examined whether a Suv39H1-regulated heterochromatin state controls migratory populations in cervical carcinomas.

Studies of cell migration have often used treatment of cells with growth factors to induce an EMT^9^. However, alternative modalities of cell migration have also been reported in solid cancers^10^, and in other contexts, EMT is not associated with migration, and is instead linked to functions such as chemoresistance^11,12^. Current EMT models may thus be insufficient in studying migration associated changes. Other studies examining metastasis have compared cells cultured from primary and metastatic tumours^13^. However, expanding these cell populations in culture may result in a loss of transient migration associated changes.

To study chromatin changes directly associated with migratory traits, we analysed migrated and non migrated populations of cervical cancer cells, functionally segregated with the help of an *in vitro* transwell set up (Fig. 1b). As invasive and pre-metastatic traits prominently emerge during the carcinoma *in situ* stage, we used the SiHa cell line, which is derived from a primary carcinoma stage. We coupled this approach with a clinicopathological analysis, using clinical biopsies and meta-analysis of RNA-Seq and clinical data from The Cancer Genome Atlas (TCGA). We observe that Suv39H1 is downregulated in migrated fractions of cells, and Suv39H1 knockdown enhances migration *in vitro*. Clinicopathological approaches revealed the emergence of a migratory Suv39H1^low^population during human cervical cancer progression, that low Suv39H1 expression correlates with lower patient survival. Finally, we observe that these Suv39H1-low migrated subpopulations are characterised by transcriptional signatures of migration, enlarged nuclei and broad scale alterations in H3K9me3 distribution.

**Figure 1.**
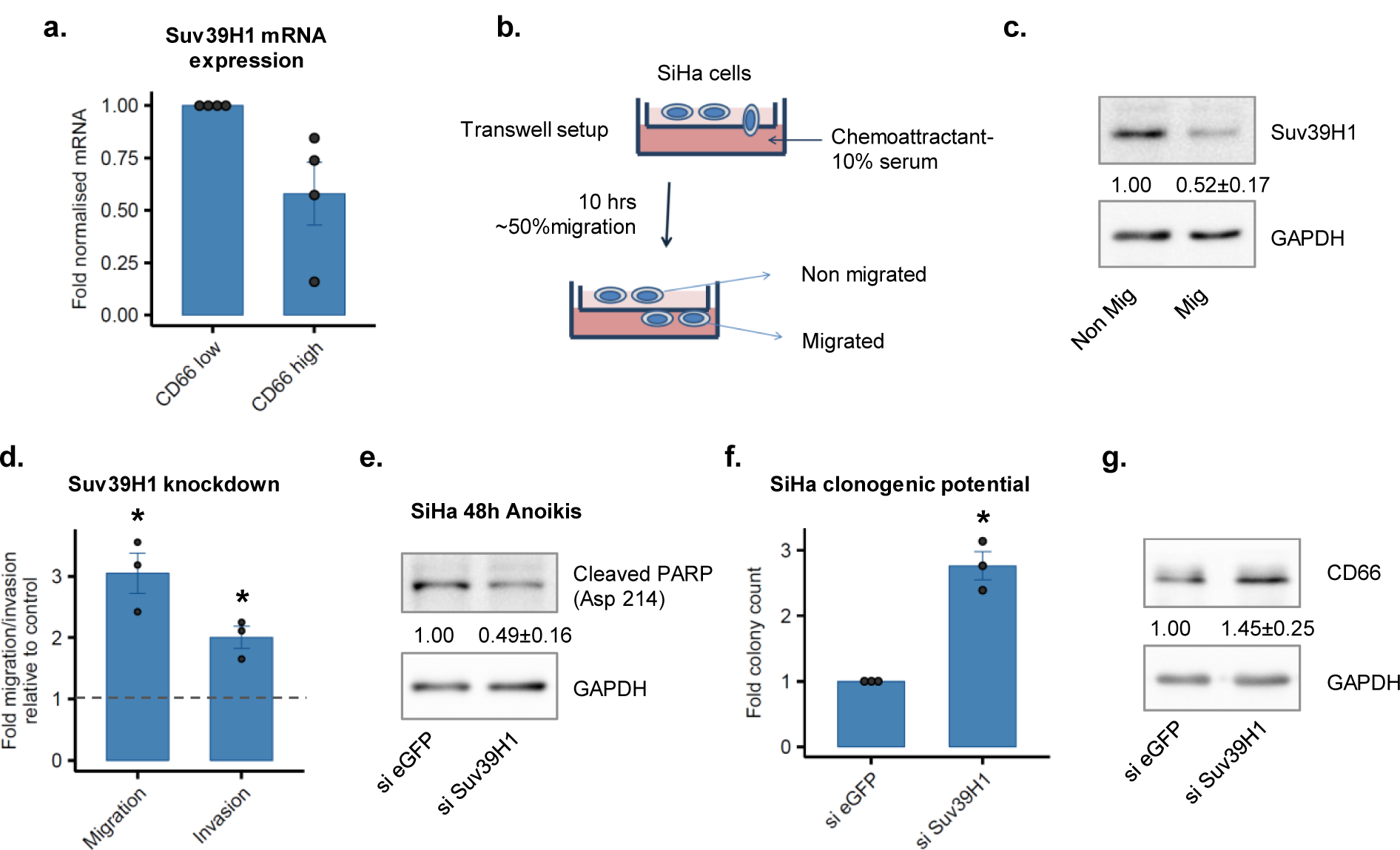
Low Suv39H1 expression characterises cervical cancer migratory states. **a.** qPCR assay showing relative mRNA expression of Suv39H1 in CD66+ve cells, enriched from CaSki spheroids (n=4) **b.** Schematic representation of an *in vitro* transwell set up, used to obtain populations of migrated and non migrated SiHa cells. **c.** Western blot showing abundance of Suv39H1 in migrated populations of cells (n=3). Mean ± S.E.M. is indicated. **d**. Quantitation of migration and invasion using a transwell assay (n=3, **\***, p<0.05). **e**. Cleaved PARP estimation by western blotting under anoikis conditions (n=4) **f.** Estimation of clonogenic potential using clonogenic assay (n=3, *, p<0.05). **g.** Western blot showing abundance of CD66 in control and Suv39H1 kd cells (n=10).

## Results

### Low Suv39H1 expression characterises cervical cancer migratory states *in vitro*

Initial observations showed a depletion of Suv39H1 in CD66 enriched spheroid cultures^1^, as well as in CD66+ve cells sorted from these cultures (Fig. 1a). Given these observations, we sought to determine whether Suv39H1 depletion is directly linked to a migratory phenotype, and whether this depletion enhances cell migration. To examine populations showing enhanced migration, we sorted migrated and non migrated populations from SiHa cells using a transwell set up. Suv39H1 protein expression was estimated in transwell derived migrated and non-migrated populations of SiHa cells, and 1.9-fold lower Suv39H1 protein was noted in migrated populations, relative to non-migrated populations (Fig. 1c). Further, Suv39H1 protein knockdown in SiHa cells using RNAi led to a 3 and 2-fold enhanced transwell migration and Matrigel invasion, respectively, and 2-fold lower cleaved PARP under anoikis conditions (Supplementary Fig. 1a, Fig. 1d, e). Under proliferation cues, these cells also show 2.7 enhanced clonogenic potential (Fig. 1f, Supplementary Fig. 1b). These phenotypes were accompanied by a 1.4-fold upregulation of CD66 at a protein level (Fig. 1g). On the other hand, overexpression of Suv39H1 resulted in a small decrease in migratory ability (1.2-fold) (Supplementary Fig. 1c, d). Thus, Suv39H1 protein is depleted in migrated populations *in vitro*, and its knockdown results in enhanced migration and invasion.

### Suv39H1^low^ cells in squamous cell carcinomas show low bulk levels of H3K9me3 and a sarcomatoid-like morphology

Following this, we sought to determine whether such Suv39H1^low^ populations are present in physiological settings, and if these putative populations show altered bulk chromatin organisation and features of enhanced migration. Current mouse models of cervical carcinogenesis do not sufficiently incorporate the physiology, variability, and patterns of metastatic spread observed in human cervical cancers^14^. To account for these aspects of cancer progression, we used a clinicopathological approach, combining histopathological analysis of human cervical biopsies with a meta-analysis of human cervical cancer data obtained from TCGA^15,16^.

Using sections from formalin fixed paraffin embedded (FFPE) cervix biopsies, we broadly categorised cells as Suv39H1^low^ and Suv39H1^high^. We also examined the bulk chromatin state of these cells, assessing bulk levels of H3K9me3, the histone mark catalysed by Suv39H1. Suv39H1^low^ regions within these sections colocalised with H3K9me3^low^ regions, in normal cervical sections as well as in Squamous Cell Carcinomas (SCCs) (Fig. 2a, b). In areas of the normal cervix showing the full spectrum of squamous differentiation, the basal and the terminally differentiated layers of the normal cervix showed cells which were Suv39H1^low^ and H3K9me3^low^, and cells in the parabasal layers were Suv39H1^high^ and H3K9me3^high^. This showed the presence of a Suv39H1^high^ progenitor population in the normal cervix. In SCC sections, cells with terminally differentiated morphology were absent. A small loss of bulk H3K9me3 expression was also noted in elongated, “sarcomatoid-like” cells in SCCs (Fig. 2b, Supplementary Fig 2). We have previously described similar sarcomatoid-like morphology in migratory CD66+ve cells in clinical sections^3^, such sarcomatoid cells have been associated with migration, metastasis and drug resistance in cancers^17–19^. These results indicated that Suv39H1^low^ cells show an association with a sarcomatoid-like morphology in SCC stages, and show lower bulk H3K9me3 levels in both normal and SCC settings.

**Figure 2.**
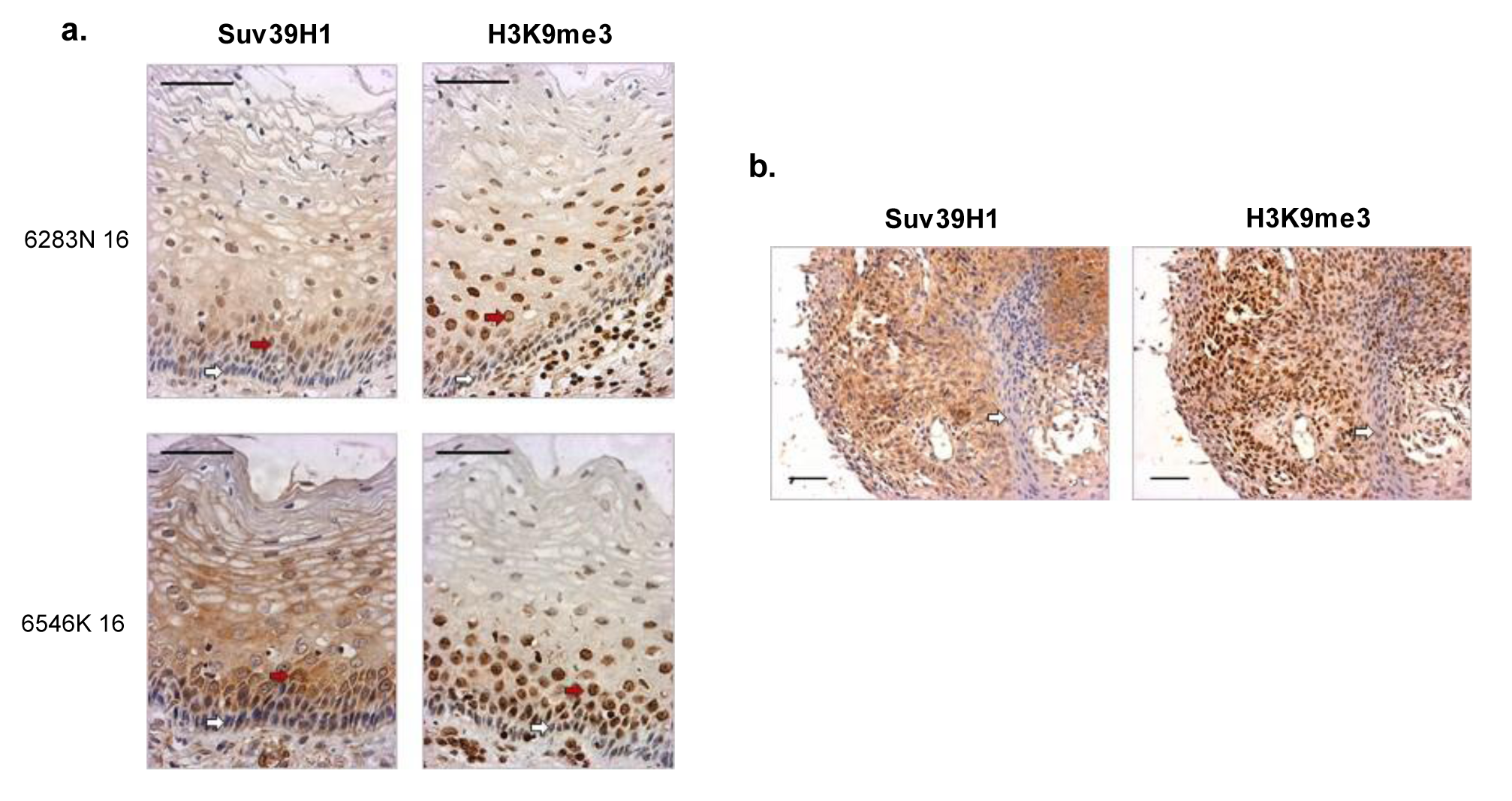
Suv39H1^low^ cells in normal cervix and in cervical carcinomas show low bulk levels of H3K9me3. Immunohistochemical staining of Suv39H1 and H3K9me3 in **a.** normal cervical epithelium (n=2, arrows indicate basal and parabasal layers) and **b.** SCC sections (n=1 of 6, arrows depict cells with sarcomatoid-like morphology). Numbers on left indicate biopsy/case numbering. Scale bar, 50 µm.

### Expansion of a proliferative Suv39H1^high^E-cadherin^high^p63^high^ cell population during early stages of cervical cancer progression

To study the emergence of migration-linked properties of Suv39H1^low/-ve^ cell populations during disease progression (Supplementary Fig. 3a), we examined p63^low/-^ E-cadherin^low/-^ regions, which mark migratory cells. p63 activity promotes keratinocyte proliferation and stemness, and its loss has been shown to favour EMT^20,21^, and E-cadherin loss is a hallmark of cell migration^22^. Some of the major phases in the progression of human cervical cancers are low grade and high grade dysplastic lesions, followed by tumours and metastasis. First, areas showing relatively normal tissue morphology within low grade squamous intraepithelial lesions (LSILs) were examined, and the presence of Suv39H1^high^p63^high^E-cad^high^ cells in the basal and parabasal layer of the epithelium was noted (Fig. 3a upper panel, Supplementary Fig. 3b, upper panel). A reduction in the frequency of Suv39H1^-ve^ cells was noted in the basal layer at this stage, relative to normal cervix sections. When areas within the same section showing LSIL morphology were examined, an expansion in the parabasal Suv39H1^high^ p63^high^ E-cad^high^ pool of cells was observed (Fig. 3a lower panel, Fig. 3b, Supplementary Fig. 3b lower panel), and no Suv39H1^-ve^ cells were noted in the basal layer. Further, the outer Suv39H1^-ve^ population with terminally differentiated morphology also showed a proportionate decrease, relative to the expanded Suv39H1^high^ population. Thus, a proliferative expansion of a Suv39H1^high^ progenitor population is noted in early stages of cervical cancer progression.

**Figure 3.**
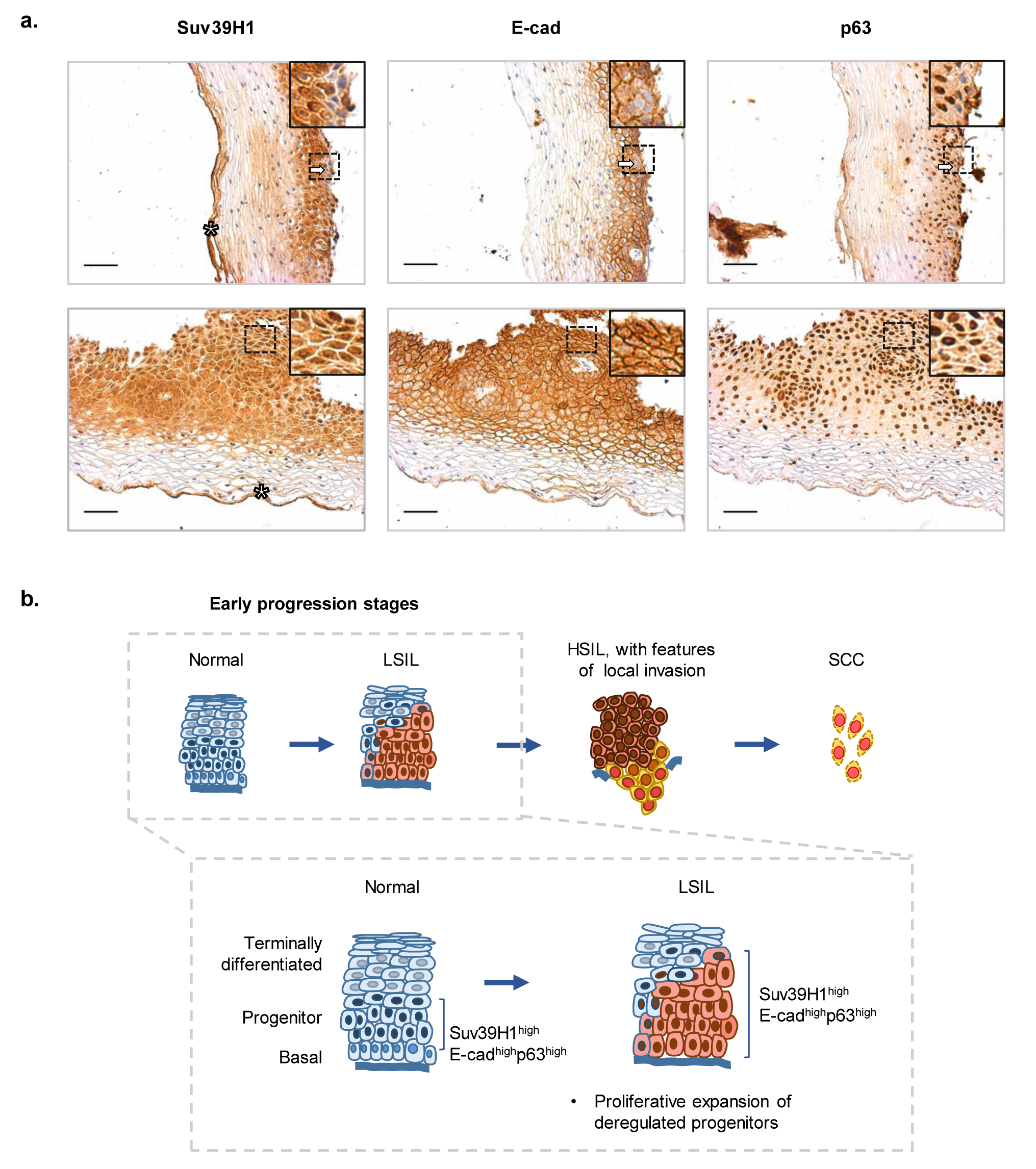
Expansion of a proliferative Suv39H1^high^ E-cadherin^high^p63^high^ cell population during early stages of cervical cancer progression. Immunohistochemical staining of Suv39H1, E-cadherin and p63 in biopsy sections. **a.** LSIL graded biopsy; Upper panel: area of section showing relatively normal morphology-arrows indicate Suv39H1^-ve^ cells and corresponding regions in serial stained sections; Lower panel: area of section showing LSIL morphology (n=1 of 2 replicates). Inset: representative enlarged regions from original images (dotted selection region). * denotes staining artefact in outermost terminally differentiated region. Scale bar, 50μm. **b.** Schematic depicting Suv39H1-linked transitions during early stages of carcinoma progression.

### Emergence of migratory Suv39H1^low^E-cadherin^low^p63^low^ cell populations during advanced stages of cervical cancer progression

We next examined sections corresponding to advanced stages of cervical cancer progression, and examined for the emergence of migratory populations. We also sought to assess modalities of migration in these putative populations, such as collective migration (characterised by a partial retention of cell-cell contacts) or single cell migration (characterised by a complete loss of cell-cell contacts). Suv39H1 distribution was first examined in a high grade squamous intraepithelial lesion (HSIL) biopsy. At this stage, the pool of Suv39H1^high^ p63^high^ E-cad^high^ cells expanded to fill the entire epithelial layer (Fig. 4a), and outer cells with terminally differentiated morphology were not observed at this stage. Further, the presence of “tongues” of epithelial cells were noted, projecting into the stromal area, suggesting early invasion. The tongues showed the presence of Suv39H1^low^ cells at the stromal front, suggesting a re-emergence of Suv39H1^low^ cells, with a non-differentiated morphology, from the Suv39H1^high^ pool. Cells within these regions were Ecad^low^ and p63^-^, suggestive of collective cell migration with a partial loss of cell-cell contact. Next, squamous cell carcinomas were examined. Here, 2 out of 5 biopsies showed pools of Suv39H1^-ve^, Ecad^-ve^, and p63^-ve^ cells, suggestive of a single cell mode of migration (Fig. 4b, Supplementary Fig.4). Other SCC sections showed less drastic losses of Suv39H1, nonetheless, pools of cells showing lower Suv39H1 also showed lower E-cadherin and p63. Thus, we note the emergence of Suv39H1 low cells in two key transitions (Fig. 4c). The first is from high grade dysplasias to invasive tumours, the emergence of Suv39H1^low^ cells from expanded pools of proliferating Suv39H1^high^ cells. The second is in the emergence of Suv39H1^-ve^ cells as potential metastatic precursors in the SCC stage.

**Figure 4.**
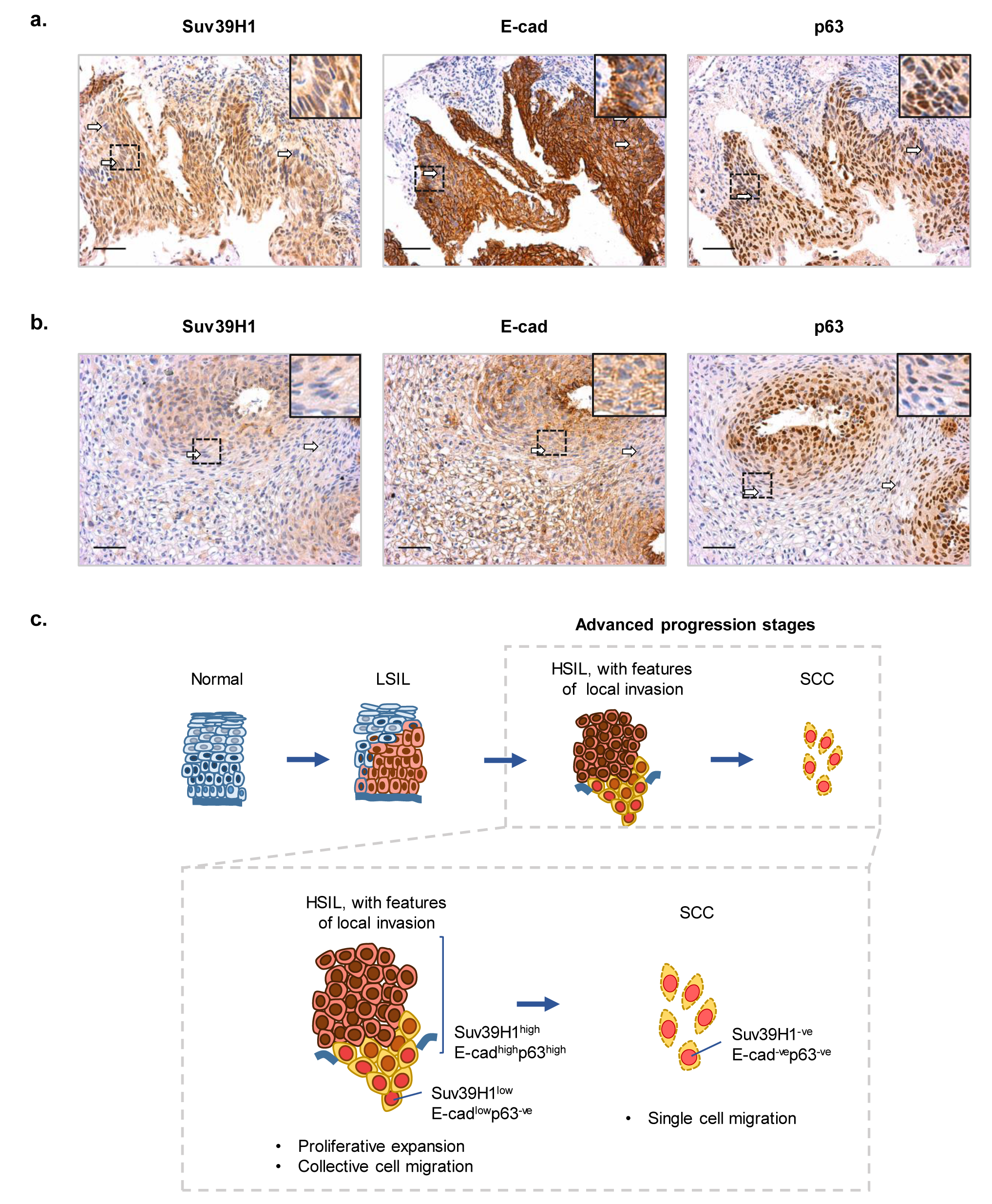
Emergence of migratory Suv39H1^low^E-cadherin^low^p63^low^ cell populations during advanced stages of cervical cancer progression. Immunohistochemical staining of Suv39H1, E-cadherin and p63 in biopsy sections. **a.** HSIL graded biopsy (n=1). Arrows indicate cells with features of migration, at areas of projection into stroma. **b.** SCC graded biopsy (n=1 of 2 showing similar trend, total n=6, ref. Supplementary data Fig. 4). Inset: representative enlarged regions from original images (dotted selection region). Arrows indicate cells with features of migration, also showing Suv39H1^low^ or Suv39H1^-ve^ cells and corresponding regions in serial stained sections. Scale bar, 50μm. **c.** Schematic depicting Suv39H1-linked transitions during advanced stages of carcinoma progression.

### Low Suv39H1 abundance in cervical carcinomas correlates with poor clinical outcomes, and with gene expression signatures of migration

Since we observed variation in Suv39H1 distribution across SCCs, we next asked if Suv39H1 abundance in tumours correlated with disease outcomes in human patients. Further, we sought to determine whether patients with low tumoural Suv39H1 showed gene expression signatures of cell migration, migration associated signalling pathways, and an association with CD66 expression signatures. We examined publicly available RNA-seq profiling data from primary human cervical tumours (Cervical squamous cell carcinoma and endocervical adenocarcinoma, CESC^15,16^), generated by The Cancer Genome Atlas (TCGA) project. Squamous Cell Carcinoma data sorted from these datasets was examined. Next, patients were sorted into Suv39H1-low and Suv39H1-high groups, based on tumoural Suv39H1 mRNA expression level. Suv39H1-low patients showed significantly lower survival over time, compared to Suv39H1-high patients (p= 0.012) (Fig. 5a, Supplementary Fig. 5a). This effect was not observed with Suv39H2, and HP1a homolog CBX5, two proteins known to synergise with Suv39H1 in some contexts (Supplementary Fig. 5b, c). Next, genes differentially expressed in Suv39H1-low tumours were examined. Consistent with the role of Suv39H1 as a transcriptional repressor, 1710 genes were upregulated and 721 genes were downregulated in Suv39H1-low tumours. These Suv39H-low tumours showed an upregulation (>1.5 fold) of several migration and matrix remodelling associated genes, including Twist1, Zeb1 and several MMPs (Fig. 5b). Gene Ontology (GO) and KEGG pathway analysis revealed a significant (p<0.01) enrichment for terms including cell migration, vasculature development, TGF-β signalling, and cell adhesion in the Suv39H1-low patient group (Fig. 5c, Supplementary Fig. 6a). Active TGF-β signalling has been reported in CD66+ve cells, and has been demonstrated to drive a switch to a non-proliferative, migratory CD49f-ve, CD66+ve cell state^3^. Further, genes upregulated in Suv39H1-low tumours show a statistically significant overlap with a CD66+ve cell profile^2^ (Fig. 5d). The Suv39H1-low group also showed a significant (p<0.01) downregulation of DNA replication and cell cycle signatures (Supplementary Fig. 6b). Thus, low Suv39H1 mRNA abundance in human tumours correlates with poor disease outcomes. Further, gene expression in these Suv39H1-low tumours shows links with cell migration, TGF-β signalling, and with a CD66+ve gene signature.

**Figure 5.**
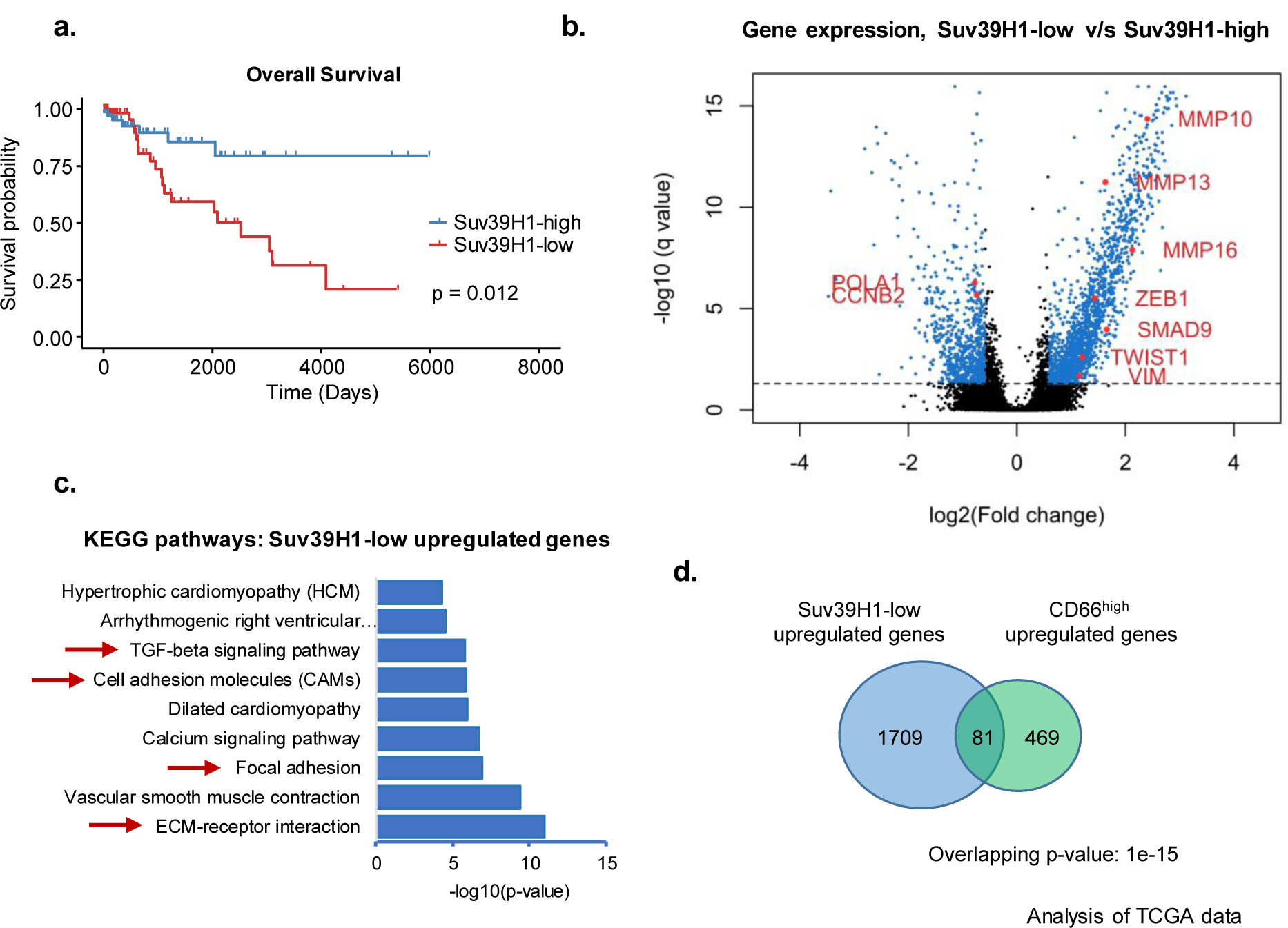
Low Suv39H1 abundance correlates with poor clinical outcomes in cervical cancers. **a.** Kaplan-Meier plot depicting survival in patients with low v/s high tumoural Suv39H1 using a median group split (n=94 for each group). **b.** Volcano plot depicting differential gene expression in Suv39H1^low^ tumours v/s Suv39H1^high^ tumours. Blue points indicate genes significant at cutoff fold change (1.5 fold) and FDR corrected q value (0.05). **c.** Enriched KEGG pathway terms corresponding to genes upregulated in Suv39H1^low^ tumours. Arrows indicate adhesion and migration associated terms. **d.** Overlap of TCGA Suv39H1^low^ upregulated genes with genes upregulated in CD66^high^ populations (Pattabiraman et al., 2014), p-value computed using Fisher’s exact test. Results shown in this figure are based on Level 3 RNA seq data obtained from The Cancer Genome Atlas Project (TCGA) Research Network (http://cancergenome.nih.gov/).

**Figure 6.**
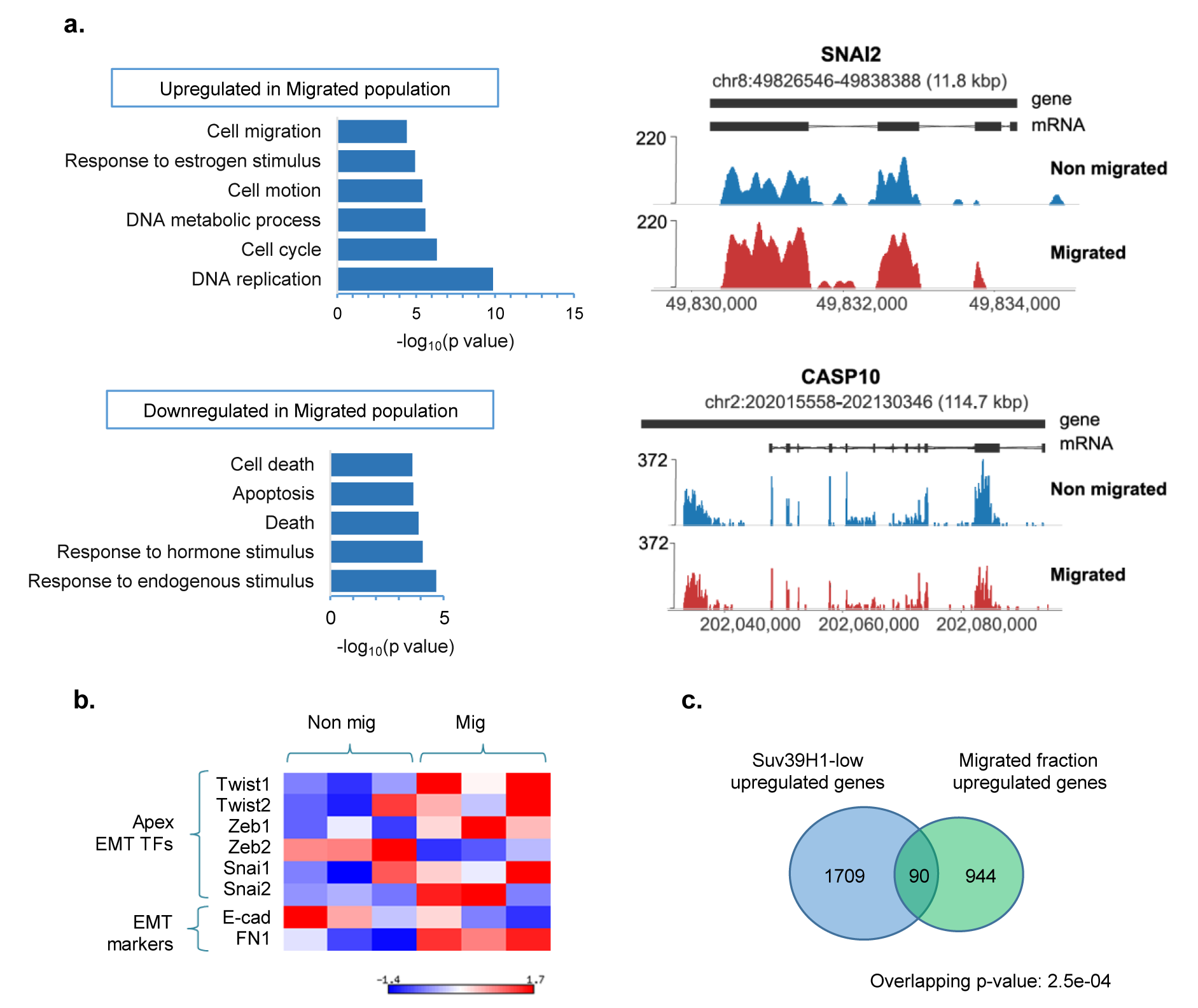
Migrated populations of cervical cancer cells show Suv39H1-linked migratory transcriptional signatures. **a.** Left, GO Biological Process terms corresponding to genes upregulated and downregulated in migrated populations (across replicates, n=3). Right, plots depicting mRNA abundance (normalised read count) of representative differentially expressed genes *Snail2* and *Casp10* in migrated and non migrated populations, in a single replicate (n=1 of 3). **b.** Expression of core EMT transcription factors and markers across replicates (n=3), depicted as a heat map. **c.** Overlap of migrated population upregulated genes with set of genes upregulated in TCGA Suv39H1-low tumours.

### Migrated subpopulations of cervical cancer cells show Suv39H1-linked changes in transcription and global H3K9me3 alterations

Given these differences in clinical outcomes and transcriptional signatures in Suv39H1^low^ tumours, we sought to determine whether a Suv39H1 and H3K9me3 low state may drive migratory populations via a chromatin-based regulation of transcription. We considered two alternative mechanisms by which Suv39H1-regulated H3K9me3 could effect putative changes in gene expression in migrated populations. One is pathway specific regulation mechanism, wherein H3K9me3 is depleted over genes within specific pathways, or on specific anti-migratory genes, such as epithelial genes^22^. This loss of H3K9me3 would allow a specific transcriptional upregulation of these genes. Another possibility is that H3K9me3 plays a broader, restrictive barrier to changes in gene expression^4–6,23^. This would entail a broader loss of H3K9me3 in migrated populations, for example, over all promoter classes. This, in turn would allow more rapid changes in transcription, effected by other, more target-specific transcription factors.

First, we sought to determine whether these migrated populations reflect migratory signatures at a transcriptional level. Transcriptional changes directly associated with migration have rarely been reported^24–26^, with a bulk of cell migration literature instead exploring post-translational signalling and cytoskeletal dynamics^10^. First the transcriptome of *in vitro* migrated cell populations was examined, by performing RNA-Sequencing of transwell-derived migrated and non-migrated populations of SiHa cells. When Gene Ontology analysis was performed on differentially expressed genes, genes upregulated in migrated populations showed enrichment for cell motion and cell migration GO terms, along with DNA replication signatures (at p<10^-4^) (Fig. 6a). Genes downregulated in migrated populations, on the other hand, showed enrichment for apoptosis and cell death (at p<10^-4^). An upregulation of known apex EMT transcription factors Twist1, Snail2, and Zeb1 was observed in the migrated population (Fig. 6b), along with a downregulation of cell-cell adhesion marker E-cadherin and an upregulation of Fibronectin. Differential expression of a small number of long noncoding RNAs was also detected in the migrated population (Supplementary Fig.7b). We also sought to assess which transcription factors effect these transcriptional changes in migrated populations, by examining for overrepresented transcription factor binding sites (TFBS) in migrated population upregulated genes. Transcription factors such as AP1, TGIF, STAT3, and SOX9 showed overrepresented target genes in migrated populations (Supplementary Fig. 7c). Further, a significant overlap was noted between genes upregulated in migrated populations and genes upregulated in TCGA Suv39H1 low patient tumours (p=2.5x10^-4^), indicating a link with a Suv39H1 transcriptional signature (Fig. 6c). Differential expression of Suv39H1 itself was not observed at the transcription level, indicating that altered levels of Suv39H1 in SiHa migrated populations are regulated post transcriptionally.

Next, bulk levels of H3K9me3 were a-ssessed by immunofluorescence and western blotting. These approaches revealed 1.6 and 3-fold reductions in H3K9me3, respectively, in migrated populations (Fig. 7a, c, Supplementary Fig. 8a, b). The nuclei in migrated populations also appeared 1.2-fold larger (Fig. 7 b, c, Supplementary Fig. 8c), suggesting a decondensed chromatin state. On the other hand, another histone mark, H3K4me3 (which is associated with transcriptional activation), did not show detectable bulk differences (Supplementary Fig. 8d-f).

**Figure 7.**
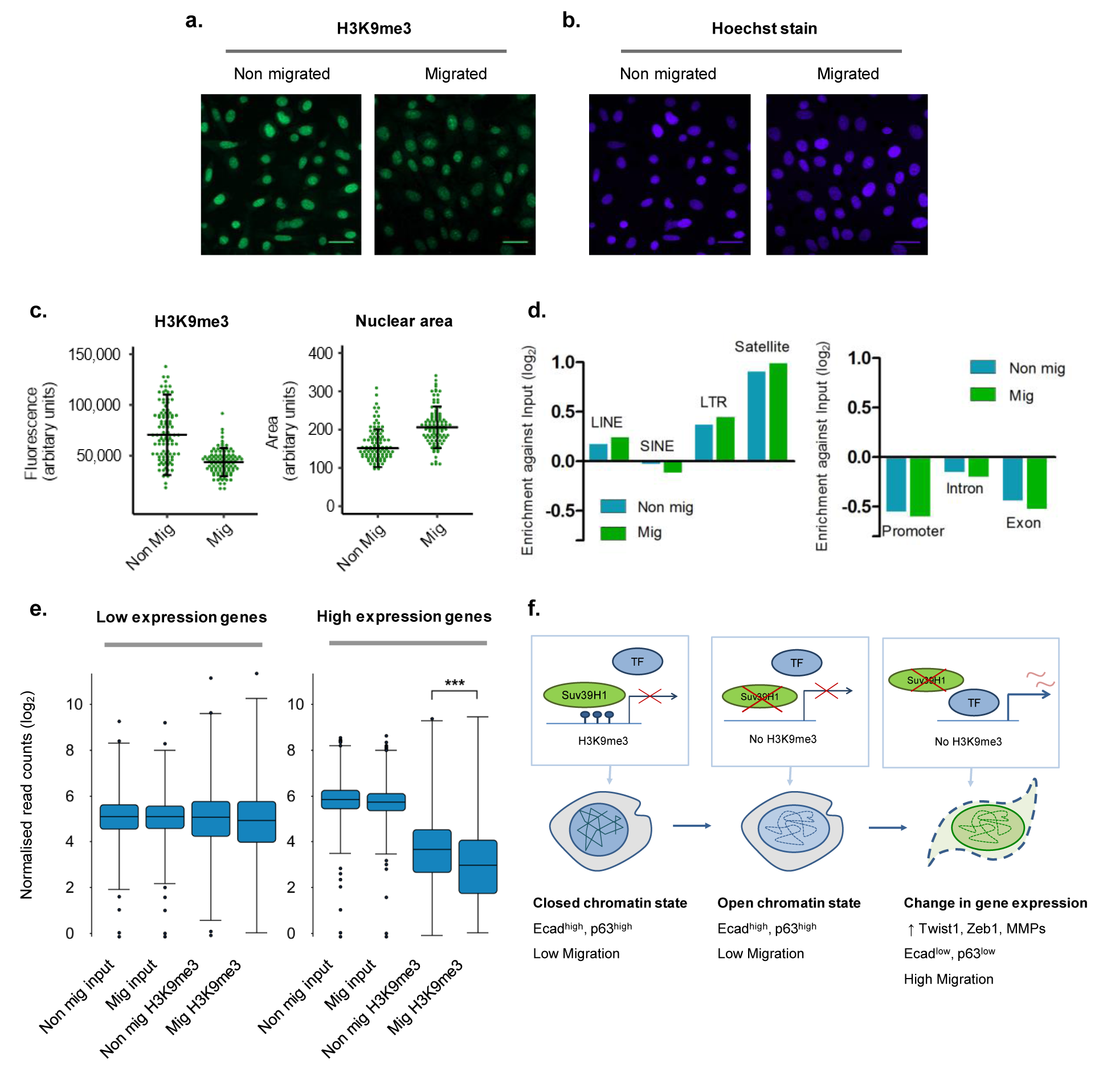
Migrated populations of cervical cancer cells show global alterations in H3K9me3 distribution. **a, b.** Representative images of H3K9me3 expression and nuclear size assessed by immunofluorescence (n=1 of 4, and n=1 of 3 replicates, respectively). Scale bar, 35μm. **c.** Quantitation of H3K9me3 staining intensity (left) and nuclei area (right) in cell populations. Bars depict median and interquartile range from a single representative experiment. **d.** Enrichment of H3K9me3 ChIP reads against input across repetitive element (left) and gene element (right) classes (n=1 of 2 replicates). **e.** Distribution of H3K9me3 ChIP reads over promoters of low expressed (0-25 percentile) and high expressed (75-100 percentile) genes (n=1 of 2 replicates),**\*\*\***, p<0.0001, Wilcoxon signed rank test. **f.** Model of Suv39H1 and H3K9me3 based regulation of cervical cancer migratory populations.

We sought to determine whether chromatin alterations in these cells acted on a locus specific, gene specific manner^22^, or whether broader chromatin changes operated across genomic classes^6,23^. Using H3K9me3 ChIP-Seq, we assessed genomic locations at which differential H3K9me3 binding is noted. H3K9me3 ChIP-seq profiles revealed broad regions of enrichment across the genome. Examining across genomic classes, an enrichment of H3K9me3 was observed over intergenic regions and repetitive elements, along with a depletion of H3K9me3 over more open chromatin- for example, exons and most promoters (Fig. 7d, Supplementary Fig. 9a). These findings are consistent with H3K9me3 binding patterns reported previously^27,28^. Further, migrated populations showed greater loss of H3K9me3 over open chromatin regions. Analysis of H3K9me3 distribution over gene bodies showed a depletion of over transcription start sites (Supplementary Fig. 9b, c), and an increase in level along the gene body, which has been previously reported^29^ and suggested to prevent aberrant transcription initiation within the gene body. Next, results of the H3K9me3 ChIP-seq were compared against RNA-seq gene expression data. No direct correlation was observed between H3K9me3 occupancy and differential gene expression in migrated and non migrated populations (Supplementary Fig. 9d). However, a correlation was noted between changes in this methylation and the absolute level of mRNA expression. While promoters of low expression genes (0-25 percentile expression) did not show significant changes in H3K9me3, promoters of high expression genes (75-100 percentile expression) showed low H3K9me3, and this mark is further depleted in migrated populations (p<10^-4^) (Fig. 7e, Supplementary Fig. 9e). Thus, migrated populations of cervical cancer cells show Suv39H1-linked transcriptional alterations, enlarged nuclei, and a broad loss of H3K9me3 over the promoters of highly expressed genes.

## Discussion

In summary, we report that a Suv39H1^low^ heterochromatin state drives migratory populations in cervical cancer. These migrated subpopulations show Suv39H1-linked transcriptional signatures of migration, enlarged nuclei and broad alterations in H3K9me3 distribution (Fig. 6, Fig. 7a-e, Supplementary Fig. 8a-c). We report for the first time, to our knowledge, an anti-migratory role for Suv39H1 in a human cancer. Suv39H1 has been suggested in roles as an oncogene, notably as a pro-migratory driver in colon and breast cancer settings, as well as a tumour suppressor, depending on the context^30–33^. This is likely because, like most chromatin regulators, Suv39H1 does not possess sequence specificity- the genomic loci to which it is recruited to depends on the transcription factor it associates with^22^, and on the chromatin landscape, and these factors may be dependent on the tissue of origin and mutation spectrum in these cancers.

We also note an association of these Suv39H1-low populations with CD66+ve populations in phenotypes, markers and transcriptional signatures. Suv39H1 knock down induces invasive and migratory phenotypes *in vitro*, and increases CD66 expression (Fig. 1d, f). Suv39H1-low tumours also show gene expression signatures of migration, an overlap with a CD66 signature, TGF-β signalling, and a depletion of proliferation signals (Fig. 5, Supplementary Fig. 6 a, b). In line with patterns observed with CD66^high^ populations, we note Suv39H1^low^ cells in advanced stages of clinical progression show sarcomatoid-like morphology, features of migration and a loss of proliferation signals (Fig. 2b, Fig. 4a, b). Lastly, similar to CD66+ve populations, we observe reflections of EMT at a transcriptional level in these migrated populations and Suv39H1^low^ tumours (Fig. 5b, c, Fig. 6b). This includes the differential expression of core transcription factors *Twist1* and *Zeb1*, MMPs, and EMT-associated cell surface markers.

One noteworthy deviation from the EMT model observed in these functionally sorted migrated cell populations is the broad depletion of H3K9me3, especially over gene promoter elements (Fig. 7a, c, e, Supp. Fig. 8 a, b). This is in contrast to the more specific loss of H3K9me3 at epithelial gene promoters observed in EMT models^22^. Together with our observations with in vitro phenotypes, gene expression signatures, and imaging, these suggest a model of increased epigenome plasticity^23^ in migrated populations- that a depletion of Suv39H1 and H3K9me3 allows genes to be more susceptible to remodelling and changes in gene expression (Fig. 7f). This would allow cells to transition more efficiently from a non migratory state to a migratory state, in the presence of environmental migration-inducing signals (e.g. TGF-β signalling). Interestingly, though the roles of Suv39H1 in cancers appear to be more context dependent, this effect of Suv39H1-mediated plasticity restriction in cervical cancers is more similar to functions reported in non-pathological processes imilar “restrictive” roles for uv and K me mediated cell fate barriers have been demonstrated in IPSC reprogramming and T-cell lineage commitment^4–8^, where loss of Suv39H1 leads to enhanced reprogramming or defective differentiation. Consistent with this notion, the basal stem cell population in normal cervix sections show a depletion of Suv39H1 and H3K9me3, relative to more differentiated parabasal populations (Fig. 2a). Plasticity-based regulation in Suv39H1 depleted cells would also explain enhanced proliferation observed under proliferation cues, and enhanced migration under migratory cues (Fig. 1d, f). Collectively, our data builds upon the notion of Suv39H1-regulated heterochromatin as a barrier to plasticity in cervical cancer cell migration, and suggests that similar plasticity may regulate other switches in cell fate.

Lastly, this study examines chromatin based changes in migratory populations with a focus on gene regulation aspects. However, it is likely that chromatin based processes may also play roles in more physical aspects of migration-such as regulating the shape and stiffness of the nucleus during invasion and migration. Further, such physical based phenomena may also be linked with gene regulation, genome stability, and cell fate. These aspects could be addressed in subsequent studies.

## Materials and methods

### Cell culture

SiHa cells purchased from A CC ere cultured in Dulbecco’s modified Eagle’s Medium DMEM with 10% FCS. Cultures were used for assays within 27 passages of acquisition from ATCC. All cultures were routinely tested for mycoplasma using Mycoalert kit (Lonza) and were consistently mycoplasma negative, and examined for retention of described morphology and growth characteristics. For transient overexpression experiments, cells were transfected with an empty vector, pcDNA3, or plX305 Suv39H1 LV V5 (DNASU plasmid repository) for Suv39H1 overexpression. Lipofectamine 2000 (Invitrogen) was used for transfection, and cells were used for molecular and phenotypic assays 48h post transfection. Knockdown experiments were performed using esiRNA, as the heterogeneous mixture of these endoribonuclease derived siRNAs provides greater target specificity than chemically synthesised siRNAs. For these experiments, mission esiRNAs (Sigma) targeting eGFP or Suv39H1 were transfected using lipofectamine RNAiMAX (Invitrogen). Cells were used for assays 48h post transfection.

### Transwell migration set up

0.5 X 10^6^ cells were counted, resuspended in DMEM containing 1% FCS and seeded into the upper well of a 24 well matrigel coated (invasion) or uncoated (migration) transwell set up. DMEM containing 10% FCS was added to the lower well of the transwell set up as a chemoattractant. Cells were allowed to migrate 10 hours at 37C after seeding. Following this, cells at the bottom of the chamber were stained with Hoechst. Cells were imaged under a Nikon Inverted Microscope ECLIPSE TE2000-S, and counted using ImageJ. For extraction of migrated and non-migrated cell fractions for western blot, RNA seq and ChIP, 0.75 x 10^6^ cells were seeded per well of a 6 well migration transwell set up. Cells were then scraped in PBS, pooled and used for further processing.

### Clonogenic potential, cell proliferation, and anoikis assays

To assess clonogenic potential, 5000 cells were seeded into wells of a 6-well plate. After 10 days, colonies were fixed using 4% PFA, and stained using crystal violet, and counted. Cell proliferation was assessed using WST1 (Roche). Cells were seeded into a 96 well plate. At different time intervals after seeding, WST1 reagent was added to wells, and absorbance readings were taken according to manufacturer’s instructions o assay anoikis, x ^4^cells were dispersed in DMEM with 1%FCS, and seeded into poly-HEMA coated plates. 48hrs later, cells were harvested and processed for estimation of cleaved PARP.

### Western blots

Immunoblots were performed using antibodies against the following: Suv39H1 (Cell Signaling Technology, D11B6), H3K9me3 (Abcam, 8898), CD66 (Abcam, ab134074), Cleaved PARP (Cell signaling technology, 9541), GAPDH (Santa Cruz Biotechnology, sc-47724). Quantitation of western blots was performed using ImageJ, using GAPDH as a control for equal loading.

### Immunohistochemistry

Formalin Fixed, Paraffin Embedded (FFPE) biopsy sections were obtained from Kidwai Cancer Institute, with appropriate institutional ethical approvals from the Kidwai Medical Ethics committee, including obtaining informed consent from each subject. Immunohistochemistry was performed on serial sections using Leica Novolink Polymer Detection system, according to manufacturer’s instructions. The following modifications were made to the manufacturers protocol. A permeabilization step was added post antigen retrieval, using 0.1% Triton in TBS (TBS-T). All washes were performed in TBS-T. Additionally, primary antibody incubation was performed overnight at 4C. Processing and imaging of each biopsy was performed individually. Thus, distribution of positively and negatively stained areas was examined within sections; comparison of absolute intensity of staining was not performed between progression stages.

### Analysis of TCGA data

Level 3 TCGA CESC RNA Seq data was obtained from the TCGA data portal (https://tcga-data.nci.nih.gov/). RNA seq RSEM gene counts as well as RSEM normalized gene counts, as well as corresponding clinical data, were downloaded. Clinical data from the samples was used to filter out adenocarcinomas, retaining only squamous cell carcinomas. Next, normalised gene count was used to divide patients based on Suv39H1 expression. Patients were ranked based on *Suv39H1* expression, and then divided using a median, tertile or quartile split. Overall patient survival corresponding to the two groups split by median was used to create Kaplan Meier plots depicted. Similar results were seen when different group splitting methods (median, tertile, quartile) were used. For RNA seq differential expression, RSEM gene counts (not normalized) corresponding to Suv39H1-low and Suv39H1-high tumours (using quartile group split) were used. Normalisation and differential expression analysis was performed using EBseq (v1.10.0)^34,35^. Genes showing a ϡ.5-fold change and a FDR cutoff of 0.05 were considered to be significantly differentially expressed, and are depicted in a volcano plot. Lists of differentially expressed genes were analysed using DAVID (v6.7) to obtain significantly enriched GO (GO BP-ALL) and KEGG terms. Overlap of gene sets and significance testing was performed using the R Bioconductor package GeneOverlap(v1.6.0)^36^, p value is computed using Fisher’s exact test

### Immunofluorescence

For immunofluorescence, transwell membranes were fixed in 4% PFA, permeabilised with 0.2% Triton X-100, blocked with blocking solution (10% serum, 0.2 % Triton X-100), stained with primary antibody (H3K9me3, Abcam ab8898, for 1 hour) followed by secondary antibody (45 minutes) at room temperature. Transwell membranes were cut out from transwells and mounted with antifade mounting medium containing DAPI (Thermo Fisher). Imaging was done on a Zeiss LSM 510 Meta confocal microscope, and image analysis was done on ImageJ.

### RT-PCR and mRNA sequencing

RNA was extracted from migrated and non migrated fractions using the Trizol method. Extracted RNA was treated with DNase, and then converted to cDNA. The cDNA was then analysed by RT-PCR, using RPLP0 as a control for equal loading. For RNA sequencing, DNAse treated RNA was used for mRNA library preparation. mRNA was purified using poly T oligo beads. Libraries were sequenced using an Illumina HiSeq 1000 using 100 bp paired end sequencing. Read quality was checked using FASTX-toolkit (v0.0.13.2), and differential expression analysis was then performed using the Tuxedo suite pipeline^37^. Reads were mapped to hg19 human genome reference using Tophat2(v2.0.12)^38^, transcript models were created from the reads using Cufflinks2(v2.2.1)^37^, and counts were scored using Cuffdiff2 (v2.2.1)^39^. Differentially expressed genes at a FDR cutoff of 0.05 were extracted with the help of the R package cummerbund (v2.16.0)^40^. Lists of differentially expressed genes were analysed using DAVID (v6.7)^41,42^ to obtain significantly enriched Gene Ontology (GO) (GO BP-ALL) and KEGG terms, as well as enrichment for UCSC ENCODE Transcription Factor Binding Sites (TFBS). Heatmap of EMT transcription factor and marker expression was generated using matrix2png^43^. Heatmap intensity denotes FPKM, rows were transformed to have mean 0, variance 1. Overlap of gene sets and significance testing was performed using the R Bioconductor package GeneOverlap(v1.6.0)^36^, p value is computed using Fisher’s exact test RNA-Seq quality control indicated high genomic coverage-383 X 10^6^ reads were generated, with㹃.22 X 10^6^ mapped reads for each RNA-Seq sample. Experiments were carried out as biological replicates (n=3), with strong correlation between replicates.

### Chromatin Immunoprecipitation (ChIP) and ChIP-Seq

To perform ChIP-Seq with low numbers of cells obtained from the *in vitro* transwell set up, H3K9me3 ChIP-Seq protocol was adapted from a low-cell histone ChIP-Seq protocol from the blueprint epigenome project (http://www.blueprint-epigenome.eu). Briefly, migrated and non migrated populations were scraped into PBS, counted, and equal cell numbers were fixed using 1% formaldehyde in PBS for 8 minutes, formaldehyde was then quenched using 10X Glycine. Cells were washed using PBS and centrifuged. Cell pellets were resuspended in lysis/ sonication buffer for 25 minutes in ice in the presence of a protease inhibitor cocktail (Roche). Sonication was then performed on a Diagenode Bioruptor water bath sonicator. Sonicated chromatin was centrifuged to clear debris, and the supernatant was diluted and added to Dynabeads Protein G (Invitrogen) conjugated to H3K9me3 antibody (Abcam, ab8898). After overnight incubation, beads were washed, then reverse crosslinked in elution buffer for 4 hours at 65C. The supernatant was then siphoned, diluted 1:2 with nuclease free water, treated with RNAse A (Sigma) and incubated for 1 hour. DNA was then purified using Qiagen MinElute PCR columns and eluted in nuclease free water. DNA was then used for qPCR and sequencing. Sequencing was performed using an Illumina HiSeq 2500 using 100 bp paired end sequencing. A high coverage was noted; a total of 793.4 million reads were generated and 729.7 million reads were mapped across 8 samples. Libraries for replicate1 were constructed using Illumina TruSeq ChIP Sample Prep kit (cat no 15027084) with gel extraction. As a high degree of duplication was noted in this replicate (∼60-92%), likely due to DNA losses during gel extraction, library preparation for a second biological replicate (replicate2), was performed with the help of Ampure XP beads, using NEB Ultra DNA library kit (ref E7370L). This resulted in much lower duplication rates (∼10-17%). Apart from library preparation, biological replicate processing was identical in all other respects. Bioinformatic analysis was performed on replicate2, and major findings were confirmed in replicate1 after sequence deduplication during analysis, using SeqMonk (v1.37). Findings from both replicates agreed with each other, with our RNA-Seq data set (levels of H3K9me3 are lower on high expression genes), and consistent with previously published reports^27,28,29^. For analysis, read quality was assessed using FASTQC software (v 0.11.5). Reads were aligned to the hg19 human genome reference using Bowtie2(v2.2.9)^44^. Aligned reads were imported into SeqMonk^45^ (v 1.37.1)for analysis of read density across genomic classes and generation of probe trend plot across genes (±2kb). For assessing H3K9me3 read distribution over low and high expression genes, RNA seq data was used to identify genes with transcription levels between 0-25 percentile and 75-100 percentile. ChIP read density was assessed around promoters of genes (defined as the region upstream of genes, from -2000 bp to +100 bp) from these two groups.

### Statistics

Bar graphs indicate mean and standard error across biological replicates n≥, unless other ise indicated. n in all figure legends corresponds to biological replicates. Significance was assessed using two-tailed t tests, and a p value of ≤ as considered as statistically significant. For immunofluorescence intensity and nuclear area quantitation, bars indicate median and interquartile range within a single experiment; two-tailed t-tests for these experiments were performed on medians from n≥ biological replicates For Kaplan Meier plots, statistical significance was assessed using a log rank test. Plots were generated using R through R Studio (v1.1.383).

### Data availability

The datasets generated and analysed in this study are included within the manuscript (and Supplementary) or have been deposited in the Gene Expression Omnibus (GEO, for RNA-Seq and ChIP-Seq data) under accession number GSE103794. All other data are available upon reasonable request.

## Acknowledgements

The authors thank Simon Andrews and Felix Krueger (Babraham Institute) for help and guidance with ChIP Seq analysis; Tapas Kundu, Simon Andrews, Colin Jamora, Aswin S.N.Seshasayee for insights and discussions; C-CAMP and Agrigenome labs for Illumina sequencing.

## Author contributions

C. R., S. K., P. V-W, C. P., and D. N. conceived and designed study plan and experiments. C. R. and C. P. Developed methodology and acquired data. C. R., S.M.N., R.V. K. performed clinical biopsy procurement, processing, and analysis. C. R., S. K., P. V-W., D. N. analysed and interpreted data. C. R., and S. K. wrote the manuscript. S. K, P. V-W., D. N., R.V. K. supervised the study.

